# Introduced trout hinder the recovery of native fish following an extreme flood disturbance

**DOI:** 10.1101/2024.09.05.611377

**Authors:** Rory S. Lennox, Angus R. McIntosh, Hao Ran Lai, Daniel B Stouffer, Nixie C Boddy, Christian Zammit, Jonathan D. Tonkin

**Author notes:** **Corresponding author:** Jonathan D. Tonkin.

## Abstract

1. In rivers, we are seeing a shift away from natural flow regimes towards larger and more frequent extreme drought and flood events. However, it is unclear how increasing intensity and frequency of extreme flow disturbances will play out alongside existing biotic pressures, such as biological invasions, to impact aquatic biodiversity. In New Zealand, vulnerable native non-diadromous galaxiid fishes face pressure from introduced trout through interspecific competition and predation, which may influence the recovery of native galaxiids after flood disturbances.
2. Here, we employed a capture-mark-recapture study across 12 sites, along a gradient of disturbance following a major flood event, to examine the impact of extreme flooding on the population structure of non-diadromous galaxiids (*Galaxias vulgaris* and *G. paucispondylus*), and the effect of trout presence on individual galaxiid growth rates recovering from this event.
3. We found a lower abundance of all non-diadromous galaxiid size classes under higher flood magnitudes, but smaller size classes (i.e., young-of-year and 1-2 year cohorts) were more impacted.
4. Furthermore, the presence of trout, whether at low or high abundances, reduced the individual growth of native non-diadromous galaxiids, despite interspecific effects being a weaker regulator of individual growth compared to conspecific effects. Moreover, trout effects on galaxiids varied by both galaxiid size and density, such that growth of smaller individual galaxiids in low densities were most affected by the presence of trout regardless of trout density.
5. In summary, our results demonstrate that non-diadromous galaxiid population dynamics in future are likely to be affected by flood disturbance regimes and introduced trout presence, the outcome of which involves a complex balance between reduced population persistence and increased individual resistance of larger individuals.
6. Conservation efforts that focus on maintaining strategically placed trout-free source populations of adult galaxiids could therefore be an important tool to enable native dispersal into trout-affected habitat and maintain population resilience in the face of increasingly larger and more frequent extreme events, given that recruitment of non-diadromous galaxiids is higher in the absence of trout.

## Introduction

The magnitude and frequency of extreme events are rapidly increasing globally (IPCC, 2021), challenging our ability to anticipate the impacts of climate change (Macinnis-Ng et al., 2024). Natural flow regimes are critical for shaping the life histories of species ranging from aquatic insects to riparian plants, but the increasing frequency and intensity of flood events are driving changes in abundance, composition, and demographic rates of aquatic communities (Poff & Zimmerman, 2010; Tonkin, 2022). Specifically, flood disturbance can directly reduce the abundance of fish through displacement-related mortality and destruction of eggs, and indirectly through a loss of habitat and resources, which reduces a system’s carrying capacity (Elwood & Waters, 1969; George et al., 2015). However, it is unclear how increasing intensity and frequency of extreme flow disturbances will play out alongside existing biotic pressures, such as biological invasions, to impact aquatic biodiversity (McIntosh & Barrett, 2022).

Globally, biotic interactions with non-native species have been credited as one of the leading drivers of declining endemic freshwater assemblages, with ecological impacts ranging from behavioural changes and transformation of food webs to the extirpation of native species (Gallardo et al., 2016; MacNeil et al., 2024; Olden et al., 2022). On an individual level, invader-driven reductions in individual growth through interspecific competition can impair the recovery of native populations via a number of different mechanisms (Healy et al., 2022). For instance, displacement from preferred habitats can negatively affect population dynamics of native fish by reducing feeding efficiency (Mills et al., 2004). In turn, slower growing individuals that take longer to reach reproductive adult stages can have lower reproductive success (Allan et al., 2021), which can limit population growth at low densities. Furthermore, small-bodied individuals tend to be more vulnerable to predation from larger individuals, therefore slower growing individuals within flood-affected populations may suffer from reduced survival to adult stages (Gascho Landis et al., 2011; Houde, 1994). Thus, strong interspecific interactions with invaders that reduce the growth of natives can have major implications for the dynamics and structure of native populations, particularly in the face of larger, more frequent extreme floods.

In New Zealand, native non-diadromous galaxiid fishes face pressure from introduced trout through interspecific competition and predation (Jones & Closs, 2017; McIntosh et al., 2010), which may influence the recovery of native galaxiids after floods. Given their dietary overlap and shared habitat preferences, strong foraging competition with trout is known to restrict native galaxiids to less preferred foraging positions with slower current, yielding lower feeding success (McHugh et al., 2012) and likely reducing individual growth. Reduced growth of individual galaxiids could have major implications for population recovery dynamics after extreme events, because growth rates control the timing of transition between ontogenetic stages and in turn the demography of populations (Houde, 1994). However, the impact of trout on galaxiid growth may not be completely negative. Previous research showed that, after accounting for total fish biomass in a reach, galaxiids grew slower in trout-free treatments than those with trout (Howard, 2007), likely due to an increase in conspecific competition. Thus, the relative strength of trout interactions in controlling galaxiid growth remains unclear, including whether trout release galaxiids from conspecific competition and subsequently modulate recovery speed.

In May 2021, the Canterbury region of Aotearoa New Zealand’s South Island experienced one of the largest floods on record for many rivers in the region (Environment Canterbury, 2021). Given the presence of previously sampled fish populations, this event presented a unique opportunity to examine the impact of extreme flood disturbances on the population structure of non-diadromous galaxiids (*Galaxias vulgaris* and *G. paucispondylus*), and the influence of trout presence on their subsequent recovery. First, we investigated size-specific abundance responses of non-diadromous galaxiids to gradients of flood disturbance. We predicted that larger galaxiids would be more resistant to increasing flood magnitudes (H1), given that larger fish are less susceptible to displacement and mortality during floods (Aedo et al., 2021; Harvey, 1987; Videler, 1993). Second, we quantified the influence of trout on individual growth rates of galaxiids recovering from varying degrees of disturbance. We expected that trout presence would reduce individual growth of non-diadromous galaxiids (H2a), particularly on vulnerable small individuals with low conspecific densities. However, given that theory predicts conspecific competition is a stronger density-dependent mechanism than interspecific (Chesson, 2000), we expected that galaxiid density would more strongly limit individual growth compared to trout density (H2b).

## Materials and methods

### Study area and general design

We sampled twelve sites across the Waimakariri and Rakaia catchments on the eastern side of the Southern Alps, Canterbury, New Zealand (Fig. 1). Each site was sampled on four occasions after the 2021 Canterbury flood event (December, January, March, April 2021-2022). Sites were selected to represent a full range of trout presence levels along a gradient of flood magnitude following the 2021 event. As such, the twelve sampled sites, which all contained non-diadromous galaxiids, were distributed into one of three trout presence categories (trout treatments: absent, rare, common) based on three decades of research (Boddy et al., 2020; Hore, 2022; Woodford & McIntosh, 2013).

**Figure 1:**
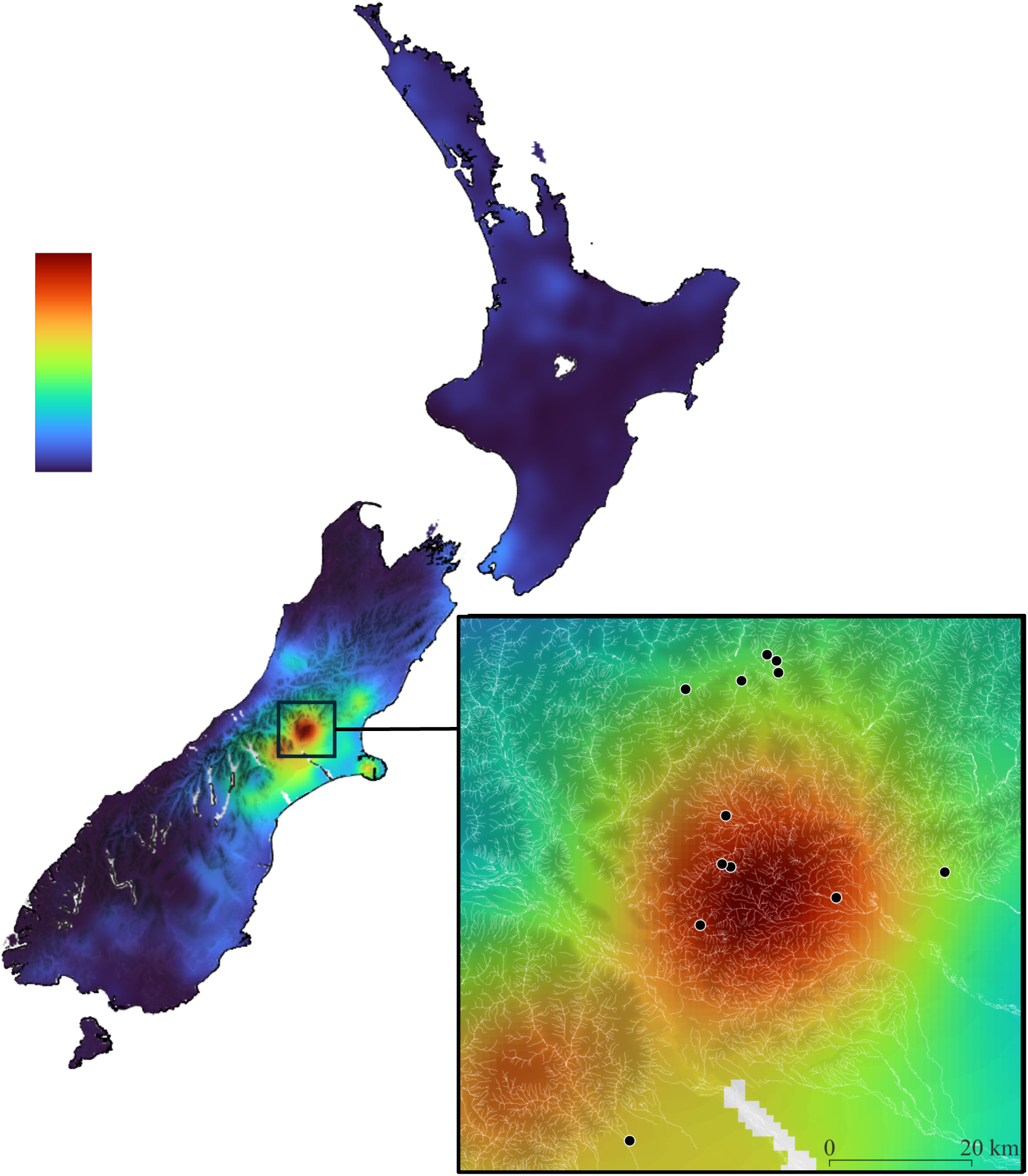
Forty-eight hour accumulated rainfall (mm) map, valid between 12:00pm 29^th^ May 2021 to 12:00pm 31^st^ May 2021, depicting the extent of rainfall that Canterbury received during 2021 flood event with the inset panel showing the epicenter of flooding in relation to the distribution of our sites (black circles). Rainfall data were supplied by Metservice New Zealand Ltd.

Assigning categorical trout treatments, instead of a continuous measure, accounted for variable detection of highly mobile trout at low density within sampling sites. At such sites, trout were categorised as ‘rare’ for the purpose of assessing trout effects on individual galaxiid growth. Of the 12 sites sampled after the 2021 flood, six sites also had pre-flood abundance estimates across three rounds during the 2020-2021 summer season (Table S1, supporting information) using the same sampling method from perennially-flowing sites (Hore, 2022). These six sites spanned a gradient of disturbance (Table S2, supporting information) that allowed for comparison, albeit with a smaller number of sites than for galaxiid growth, of pre- and post-flood galaxiid abundance.

The two species of galaxias captured during our study*, Galaxias vulgaris* and *G. paucispondylus,* were treated as a single non-diadromous galaxiid guild due to known life history and behaviour similarities (Jones & Closs, 2016). Thus, interactions among *G. vulgaris* and *G. paucispondylus* are collectively referred to as conspecific when analysing effects of flooding on population structure. However, species identity was included as a covariate within the growth model to account for known differences in maximum length between the two species. Similarly, the two species of trout captured during sampling, *Salmo trutta* and *Oncorhynchus mykiss*), are treated as a single guild in the context of interspecific effects on galaxiid growth, and collectively referred to as ‘trout’.

### Flood magnitude estimation

Most rivers within the study area are ungauged. Thus, it was not possible to measure the variation in the magnitude of the flood event directly. However, we used modelling to estimate the annual recurrence interval (range: 2- to 70-year flows) of the flood for each site, hereafter referred to as flood magnitude, using the New Zealand Convective Scale Model (NZCSM) associated with the TopNet hydrological model (Clark et al., 2008) as per the New Zealand Flood Awareness System (Cattoën et al., 2022) (for more details see Appendix 1, supporting information). Higher values of annual recurrence interval indicate flood events of higher magnitude and rarer occurrence. These values were consistent with expected magnitudes based on 27 years of observations of geomorphological changes within these streams (e.g., Boddy & McIntosh, 2017; McIntosh & Townsend, 1994; Woodford & McIntosh, 2013), and based on observed geomorphological changes after this flood event (Fig. S1, supporting information).

### Fish sampling and tagging

To estimate the abundance and size structure of non-diadromous galaxiids populations, we conducted three-pass quantitative electrofishing using a Smith-Root LR-24 Backpack Electrofisher (Smith Root Inc., Vancouver, WA, USA) following McIntosh (2000). Stop-nets were placed at the top and bottom of 30-m reaches to physically and demographically ‘close’ populations during sampling, thus we assumed that no immigration, emigration, recruitment, or mortality occurred during each sampling event.

We anaesthetised all captured fish with 20 mg/L of AQUI-S™ anaesthetic (AQUI-S New Zealand Ltd., Wellington, NZ), before recording species identity and length (to the nearest mm) for each individual fish (total length, TL, for galaxiids and fork length, FL, for salmonids). Fish were processed and recorded according to the pass they were captured on. To track individual growth, all captured galaxiids >60 mm in length were marked with a unique visual implant elastomer (VIE) tag (Northwest Marine Technology Inc., Anacortes, WA, USA) (for more details see Fig. S2, supporting information). All processed fish were placed in a bucket of cool, oxygenated stream water to recover (once swim-ability had returned), before being released back into the stream at the approximate location of capture.

To maximise recapture rates, we also conducted a single pass of electrofishing up to 50 m above and below each designated reach without the use of stop nets. Adult galaxiids (>60 mm TL) caught outside the reach were kept separate from those caught within, and were recorded as such, to be excluded from abundance and density estimates. However, these fish were tagged and measured during each sampling period to measure growth rates. Fish caught outside of the 30-m study reach were assumed to experience equivalent abiotic and biotic conditions as those caught within the reach, given the sampled reach was physically open between sampling events and the suggested *G. vulgaris* home range is ∼200 m (Cadwallader, 1976).

### Estimating size-class abundances from multi-pass sampling

To pre-process response variables for modelling of population abundances before and after the flood event, we followed previous studies (Boddy et al., 2020; Woodford & McIntosh, 2013) and arranged galaxiids into three size classes depending on individual length; young of year (YOY, < 60 mm), 1-2 year old (1-2, 60–90 mm), and ≥2 years old (2+, >90 mm). From the multi-pass depletion data, we estimated total population abundances for each galaxiid size class at each site and sampling round using a weighted maximum likelihood model (Carle & Strub, 1978), implemented with the ‘removal’ function from the *FSA* R package (Ogle, 2018). Size-class abundances at each site and sampling round were combined to calculate total galaxiid abundance and serve as a predictor variable when modelling conspecific effects on individual growth.

### Modelling population abundance

To test the effects of flood magnitude (calculated as the scaled annual recurrence interval) on galaxiid size-class abundance (i.e., H1), we modelled pre-processed counts of each size class *k* (YOY, 1-2 year old, and 2+ year old) in site *j* at each sampling round *t*, *N*_*k*j*t*_, in a generalised multi-level model as negative-binomial distributed response variables with a log link function. Consistent with other studies (Mathewson et al., 2012), we used log area (m^2^), log *A*_j*t*_, as an offset variable to account for site area that varied among sampling occasions due to fluctuations in water flow, and to allow the model to predict density (i.e., count per area).

The count model was therefore:

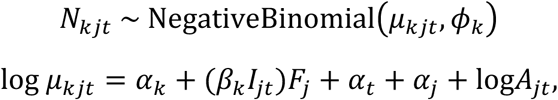

where α_*k*_ is the size-class specific intercept. The slope β_*k*_denotes the size-class specific response to flood magnitude, *F*_j_. The binary indicator variable *I*_j*t*_ equals zero for pre-flood data and equals one for post-flood data, such that *F*_j_ only affects galaxiid counts via its slope β_*k*_ when those counts were post-flood. The parameter Φ_*k*_ is the size-class specific dispersion of the negative-binomial distribution. Site and sampling round were included as random effects, α_j_ and α_*t*_ respectively, to account for any non-independence among sampling rounds across sites. Before model fitting, flood magnitude was scaled to have unit standard deviation.

After estimating the model parameters, the expected change in abundance (individuals per 100 m²) in response to increasing flood magnitude was predicted for each size class (YOY, 1-2 year, 2+ year). To do this, we predicted pre- and post-flood abundances along the flood-magnitude gradient to obtain the expected change in abundance, while marginalising over the random site and sampling-round intercepts, and setting area to 100 m².

### Modelling individual size growth

To test the effects of total galaxiid abundance and trout presence on individual galaxiid growth (i.e., H2a and H2b), we modelled with individual galaxiid lengths through time across 12 sites using the von Bertalanffy growth equation (von Bertalanffy, 1938) (for more details see Appendix 1, supporting information) in a generalised nonlinear multilevel model:

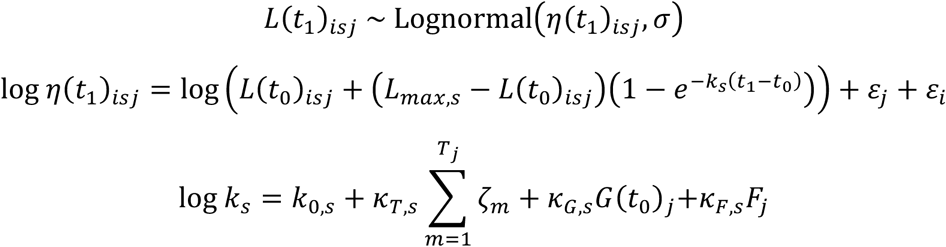

where *L*(*t*_1_)_isj_ represents the length (mm) of individual *i* of species *s* at site *j* during recapture time *t*_1_, *L*_max,s_ represents species-specific (either *G. vulgaris* or *G. paucispondylus*) maximum length, *L*(*t*_0_)_isj_ represents length of that same individual at initial capture time *t*_0_; *k*_s_represents the species-specific growth coefficient, which was allowed to be influenced by trout presence (common, rare, absent), total galaxiid abundance (scaled no. per m²), and flood magnitude (scaled annual recurrence intervals). In the *k*_s_ submodel, *k*_0,s_ denotes the species-specific intercepts; we used a sum-to-one contrast by dummy-coding *G. vulgaris* as −1 and *G. paucispondylus* as 1, so that setting the dummy species identity to zero allows us to obtain an “average” length growth prediction between species. The slope *K*_T,s_ denotes the species-specific monotonic response to trout presence T_j_ at site *j* (to ensure the effect of common trout level > rare trout level > no trout present, or common trout level < rare trout level < no trout present). The parameter ζ_m_ is a simplex parameter required to fit the monotonic trout effect (see Bürkner & Charpentier, 2020 for more information). The remaining species-specific coefficients *K*_G,s_ and *K*_*F*,s_ denote the effects of total galaxiid abundance G(*t*_0_)_j_ in the previous recapture and current flood magnitude *F*_j_, respectively. Lastly, we included the site random intercept ε_j_ to account for any non-independence of multiple recaptures from the same site over the season and environmental characteristics affecting growth, such as temperature, as well as the individual random intercept ε_i_to account for any non-independence of multiple measurements from the same individual (i.e., recaptured more than once).

The coefficients *K*_T,s_, *K*_G,s_ and *K*_*F*,s_ are not directly comparable as their corresponding predictors are a mix of continuous and ordinal variables. To facilitate interpretation, we range-standardised the predictors so they have comparable coefficients. We opted for range standardisation because the coefficient of the only ordinal predictor, trout presence, *K*_T,s_ can be interpreted as the effect across the entire range of trout levels (increasing from “absent” to “common”), as the term ∑^T$^ ζ_m_ = 1 when summing from the lowest to highest categories of trout presence. Therefore, we simply standardised the other two coefficients to be the flood effect from the lowest to highest flood magnitude, and the conspecific effect from the lowest to highest total galaxiid density, following the procedures in Lefcheck (2016).

All models were fit with Bayesian inference with Hamiltonian Monte Carlo (HMC) sampling using the *brms* package (Bürkner, 2017; version 2.18.0) and specified priors (for more details see Appendix 1, supporting information). For each model, we ran 4 HMC chains, each consisting of 5,000 total sampling iterations and 3,000 warmup iterations. Chains were checked for convergence using the Gelman-Rubin diagnostics (Ř < 1.1) and visual assessment of the trace plots. All statistical analyses were carried out in R version 4.2.1 (R Core Team, 2022).

### Growth scenarios

To enable illustration of any density-dependent effects of conspecific interactions or galaxiid size, individual galaxiid growth (length over time) was predicted from the modelling above under a range of biotic scenarios (i.e., individual galaxiid size in relation to galaxiid abundance). Specifically, we projected individual length increment in time across each level of trout presence (common, rare, and absent) under three levels of galaxiid density (fish per m²; −1 SD, average, +1 SD), and further crossed with three levels of initial capture length of individual galaxiids (mm; −1 SD, average, +1 SD) using the 419 individual galaxiids recaptured. Predictions were made with 90% credibility intervals, using counterfactual data generated for every five days of the fish sampling duration, while holding flood magnitude and galaxiid species at their averages.

## Results

Across the twelve sampled sites and over the four post-flood sampling occasions, we tagged 1,256 individual non-diadromous galaxiids and recaptured 419 fish (some more than once). Specifically, there were 158, 169 and 92 recaptures across trout absent, trout rare and trout common treatments, respectively. Additionally, we captured a total of 595 trout (mostly *Salmo trutta* and some *Oncorhynchus mykiss*), 34 longfin eels (*Anguilla dieffenbachii*), and 125 upland bullies (*Gobiomorphus breviceps*) over the four rounds of sampling.

### Density responses to extreme flood disturbance

Overall, galaxiid density varied by size classes, as indicated by different intercepts of YOY (−2.43; 90% credibility intervals or CIs: −3.49 to −1.40), 1-2 year olds (−1.36; 90% CIs: −1.88 to −0.84), and 2+ year olds (−2.64; 90% CIs: −3.17 to −2.08). These results suggest, in general, 1-2 year individuals were more common than YOY and 2+ year individuals. There were strong negative effects of increasing flood magnitude on galaxiid abundance across all sizes; flood disturbance negatively affected all galaxiid sizes, becoming more detrimental from −1.48 for YOY to −0.94 for 2+ year-old fish (Fig. 2).

**Figure 2:**
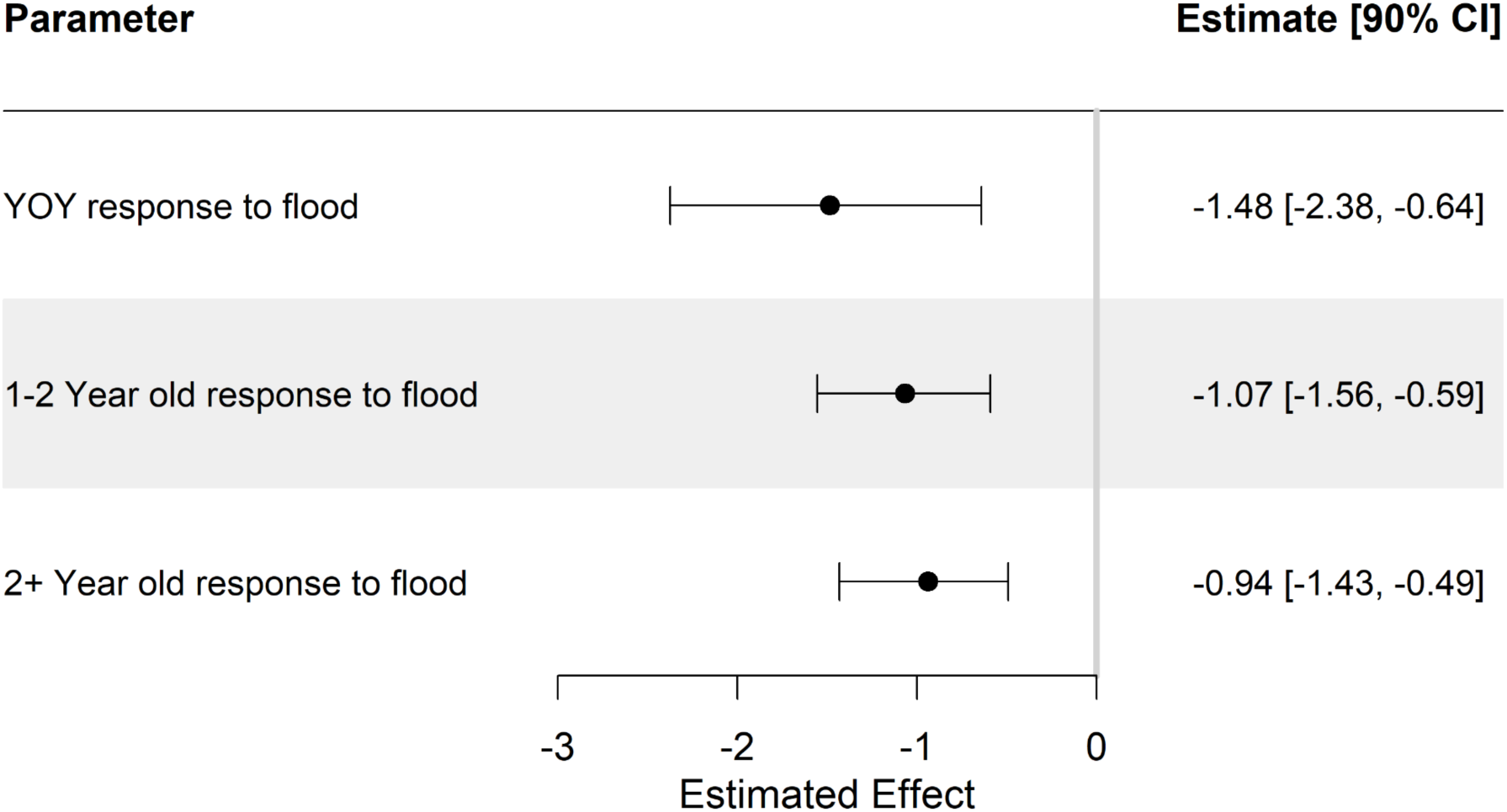
Estimated effects of flood magnitude on abundance of non-diadromous galaxiids of different sizes. Points represent posterior medians and bars indicate 90% credibility intervals.

Further comparison of galaxiid populations across the flood magnitude gradient using predicted changes in log density indicated the 2+ year size class displayed the highest resistance to increasing flood magnitude out of all three size classes (right panel, Fig. 3), in support of hypothesis 1. By contrast, the greatest negative impact of increasing flood was observed in YOY (left panel, Fig. 3).

**Figure 3:**
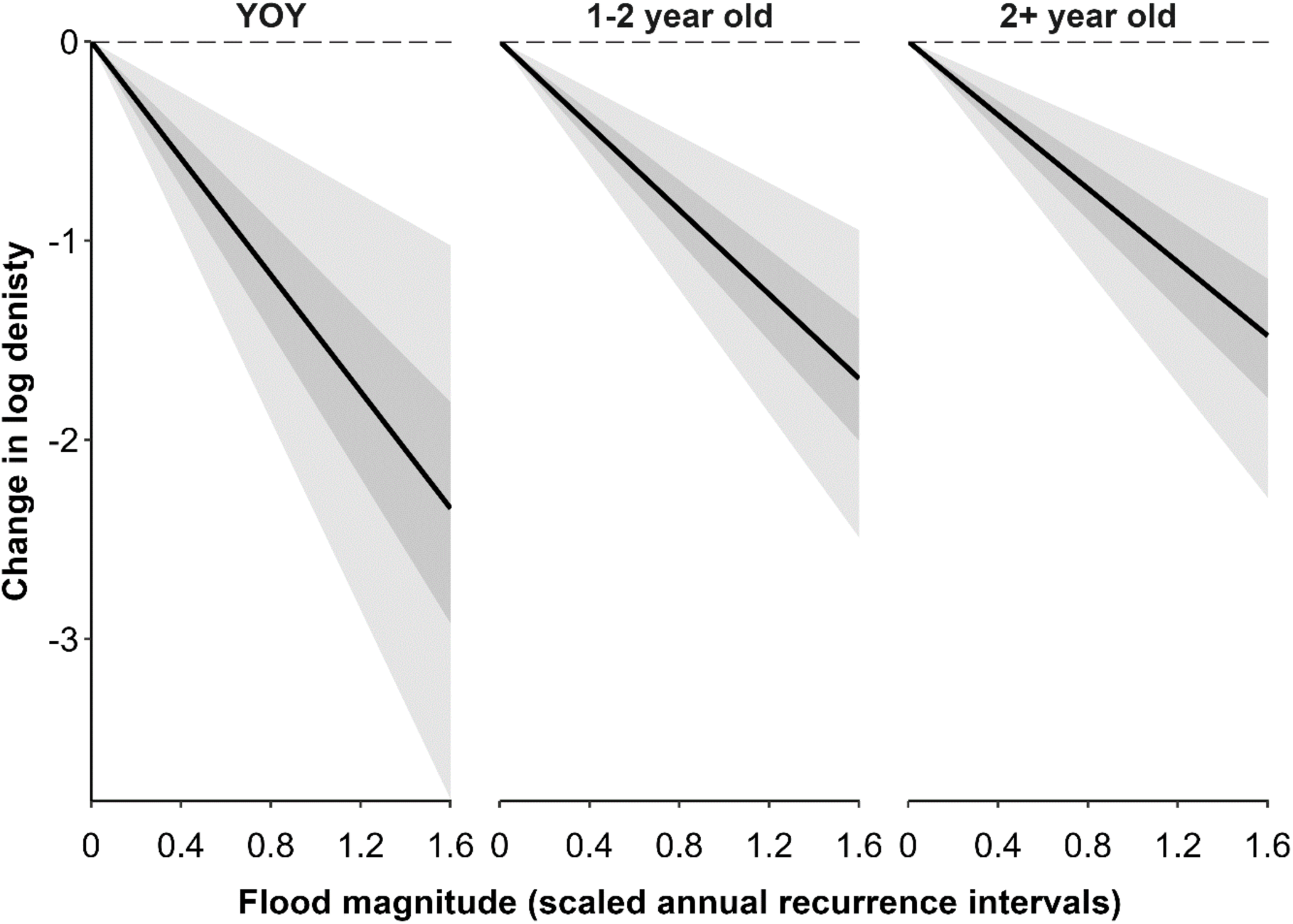
Model-expected change in log density of non-diadromous galaxiids of different sizes (YOY, < 60 mm; 1-2 year, 60–90 mm; 2+year, >90 mm) before and after the 2021 Canterbury flood in response to scaled annual recurrence intervals, generated from New Zealand Convective Scale Model (NZCSM) simulations. Larger values of flood magnitude represent larger floods, where 0 represents no flood magnitude (1-year flood) and 1.6 represents our maximum observed magnitude (70-year flood). Solid slopes represent median expected values, darker shading represents 50% credibility intervals and lighter shading represents 90% credibility intervals. Dashed horizontal lines represent no change.

### Growth responses to competition and extreme flood disturbance

Individual galaxiid growth was most strongly influenced by galaxiid abundance (range-standardised effect: −0.57; 90% CIs: −0.85 to −0.32; Fig. 4), suggesting conspecific abundance was a stronger limiter of growth than trout abundance. However, trout abundance also reduced individual galaxiid growth (range-standardised effect: −0.13; 90% CIs: −0.21 to −0.04). Finally, the standardised effect of flood magnitude was positive but weak (range-standardised effect: 0.21; 90% CIs: −0.05, 0.44).

**Figure 4:**
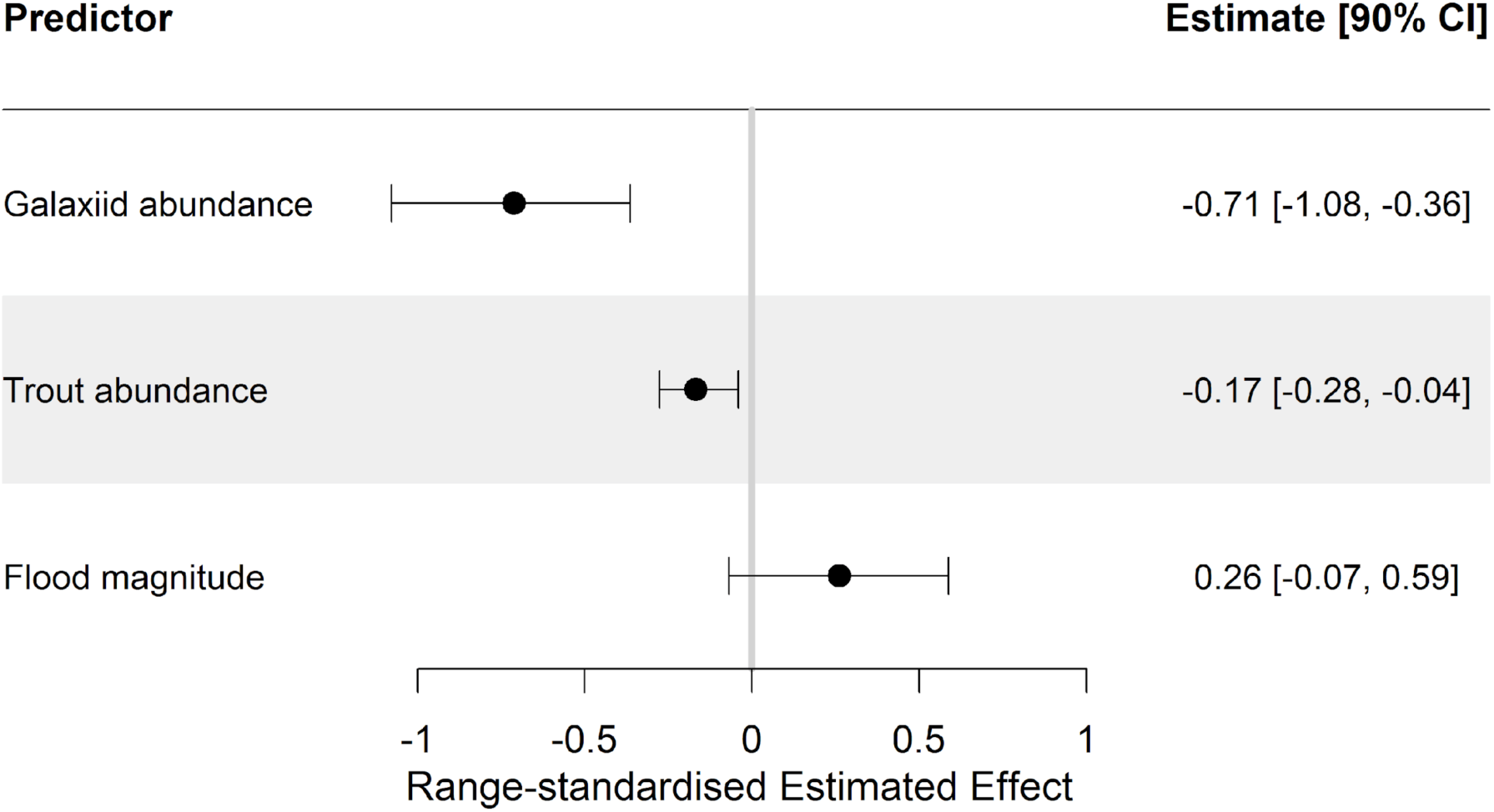
Range-standardised estimated effects of non-diadromous galaxiid abundance, trout abundance and flood magnitude on non-diadromous galaxiid individual growth (log growth coefficient, *logk*_s_). Points represent median posterior estimates and bars indicate 90% credibility intervals.

Although trout abundance had an overall negative, additive effect on individual galaxiid growth coefficient *k*_s_ regardless of initial length or galaxiid density, the effect of trout presence on individual galaxiid’s length trajectory can be non-additive due to the nonlinear nature of the von Bertalanffy growth model (Fig. 5). Specifically, trout had the greatest negative influence on individual growth of small bodied galaxiids (1 SD below average initial capture length) in low galaxiid density streams (1 SD below average galaxiid density) (Fig. 5, bottom left plot). In the absence of trout, however, small bodied galaxiids within low galaxiid density streams demonstrated the fastest growth out of all possible scenarios (Fig. 5, bottom left plot, blue line). Together, these results support the expectation that smaller bodied galaxiids at low densities would show the greatest reductions in individual growth in the presence of trout, especially compared to large-bodied galaxiids at high densities.

**Figure 5:**
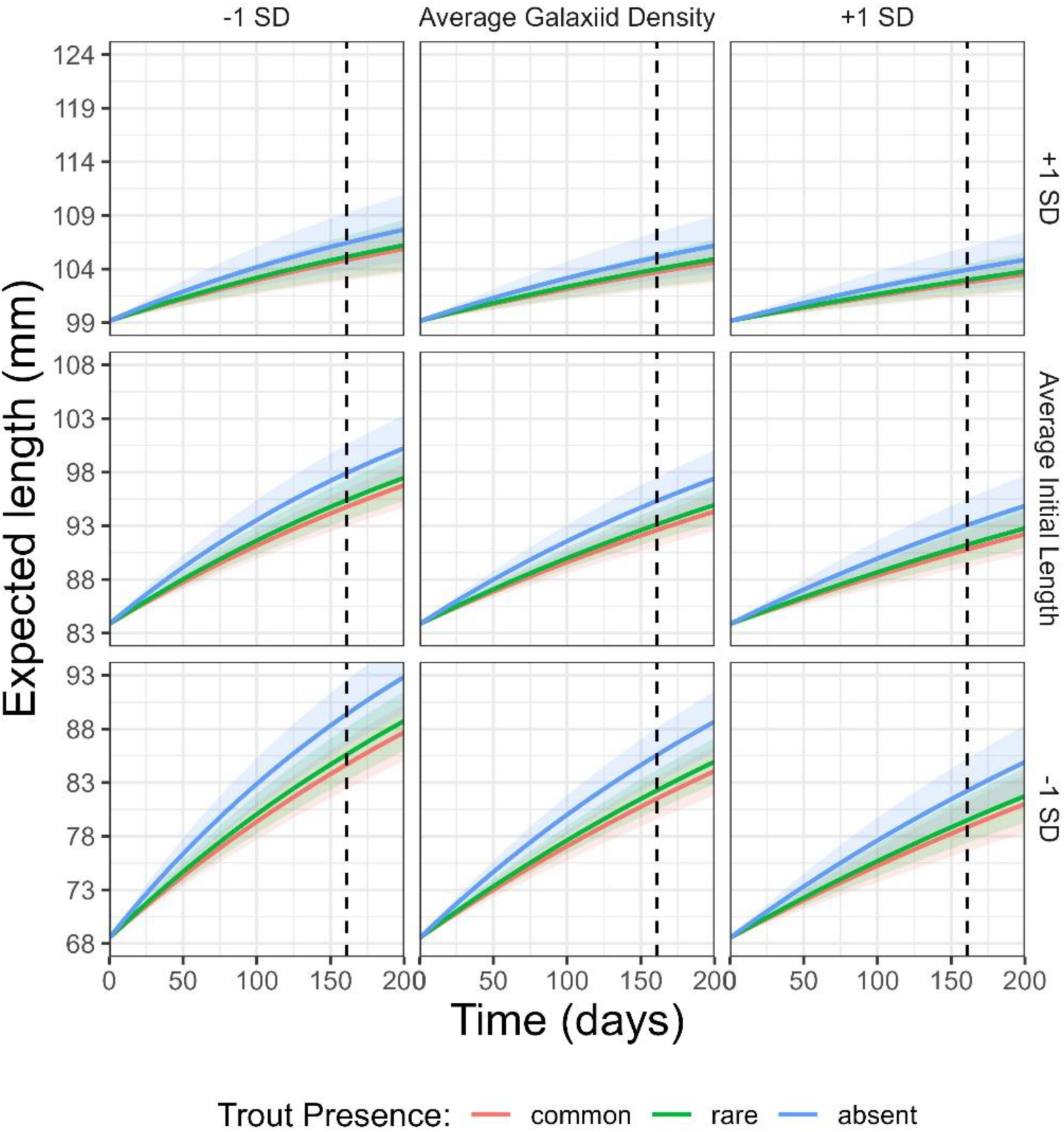
Individual growth plots predicting median non-diadromous galaxiid length (mm) through time (days), with 90% credibility intervals. Coloured lines represent trout presence levels (common, rare, absent). Time range on the X-axis is based on the mean time elapsed between recapture intervals during field sampling (53 days). Rows of plots represent variation of initial capture length of individual non-diadromous galaxiids (mm) based on observed data (average: 83.82; +/-1 standard deviation: 99.13 & 68.50). Columns of plots represent variation of non-diadromous galaxiid density (fish per m²) based on observed data (scaled average: 0.446; +/-1 standard deviation: 0.809 & 0.082). Vertical dashed lines represent the maximum observed time between individual growth measurements (161 days). We note that there are no statistical interactions between the predictors galaxiid density, initial length and trout presence in the growth model, but the model’s nonlinearity can lead to apparent “synergistic” effects.

## Discussion

Understanding how native fish populations respond to extreme disturbance and how introduced species influence their recovery is needed to inform conservation practices under changing environmental conditions (Carosi et al., 2023). Here, we examined the impact of an extreme flood on the population structure of non-diadromous galaxiids along a disturbance gradient, and the effect of trout presence on individual growth rates of galaxiids recovering from this event. We found a lower abundance of all non-diadromous galaxiid size classes under higher flood magnitudes, but larger size classes (i.e., 2+ year olds) were less impacted (supporting H1). The presence of trout, whether at low or high abundances, reduced the individual growth of native non-diadromous galaxiids (supporting H2a), despite interspecific effects being a weaker regulator of individual growth compared to conspecific (supporting H2b). Moreover, trout effects on galaxiids varied by both galaxiid size and density, such that growth of smaller individual galaxiids in low densities were most affected by the presence of trout regardless of trout density. Collectively, these results indicate that non-native trout are hindering the recovery of vulnerable populations of native species.

### Impacts on population structure

Our results revealed that non-diadromous galaxiids across all size classes were negatively affected by increasing flood magnitudes, which is consistent with an abrupt loss of biomass during extreme disturbance events (Walker, 2012). Peak flows during floods events can scour stream beds and displace entire aquatic communities, including fish, invertebrates, and algae (Death, 2008), and as the magnitude of flooding increases, so does the removal of individuals from populations (McIntosh & Barrett, 2022). Here, the gradient of flood magnitude experienced ranged from approximately annual, whereby important ecological functions are provided, to multi-decadal, whereby channel geomorphology is reset across the riverscape and widespread ecological impacts occur (Ward et al., 2002). For example, Dry Stream, a historic stronghold for native galaxiids within the Canterbury High Country, experienced a 70-year flood magnitude that resulted in extensive vertical scouring of the stream bed and a dramatic loss of galaxiids across all size classes. However, our results demonstrated that the effects were not consistent across size classes, with flood impacts known to scale with fish age (Harvey, 1987).

The negative effects of increasing flood magnitudes on non-diadromous galaxiid abundance were most pronounced within smaller size classes (i.e., YOY), consistent with hypothesis 1. Given that population connectivity can depend on dispersal of juvenile fish (Jones & Closs, 2016), extreme flood events that disproportionately remove smaller size classes (i.e. YOY and 1–2-year size) can limit recruitment and reduce connectivity between galaxiid populations on a landscape scale. This may be particularly problematic for the persistence of native populations in catchments where they interact with introduced trout, which have been recorded to reach unprecedented densities following catastrophic flood events in the USA (George et al., 2015; Roghair et al., 2002). Therefore, flood impacted populations with reduced recruitment may face an increased risk of local extirpation from connected source populations of trout.

Larger adult galaxiids were more resistant to the negative effects of increasing flood magnitude compared to younger size classes (H1), which may be explained by physiological and behavioural advantages of larger fish that make them more resistant to floods (Harvey, 1987; Kopf et al., 2014; Videler, 1993). For instance, larger fish have greater propulsion potential that makes them stronger swimmers during high flow events (Heggenes & Traaen, 1988; Jones & Closs, 2016). Furthermore, adult galaxiids have displayed refuge seeking behaviour during high flow, such as purposeful micromovements (<5 m) into low-velocity habitats (David & Closs, 2002), that may be too energetically costly for juveniles. Adult galaxiids that persist through extreme floods could in turn make up for flood-reduced recruitment through compensatory mechanisms, such as reproduction, growth, and dispersal (Ali et al., 2003; Rose et al., 2001). For example, past research has found absolute fecundity in Australia’s critically endangered Stocky galaxias (*G. tantangara*) to increase four times from 211 oocytes for a 76-mm fish to 810 oocytes for a 100-mm fish (Allan et al., 2021). Thus, flood resistant 2+ year old individuals (>90mm) are likely to be effective breeders that are important for increasing population growth at low densities and maintaining population persistence, compared to smaller individuals. Furthermore, dispersal of adult *G. vulgaris* from trout-free sites is one of the main mechanisms responsible for maintaining the persistence of native galaxiid sink patches in the presence of trout (Woodford & McIntosh, 2010). Therefore, adult galaxiid survivorship will be particularly important for compensating for reduced recruitment and may hold the key to driving recovery of flood-impacted populations.

### Impacts on individual growth

Trout abundance reduced the individual growth of non-diadromous galaxiids (H2a), across disturbance magnitudes, representative of antagonistic interactions. Specifically, introduced trout impact native fishes through inference competition in the form of fighting for prime feeding habitat and exploitative competition in the form of shared food resources (Crowl et al., 1992; McDowall, 1968; McIntosh, 2000; Townsend & Crowl, 1991; Woodford & McIntosh, 2013). Even at early life stages, trout are substantially larger and more competitive than native galaxiid species (Jones & Closs, 2016). Thus, frequent predator avoidance strategies of galaxiids that minimise antagonistic interactions with trout, like altered feeding behaviour and microhabitat use (Crow et al., 2010; Crowl et al., 1992; Glova & Sagar, 1993), could increase energy expenditure and restrict energy input. For example, Edge et al. (1993) observed *G. vulgaris* making fewer feeding attempts in the presence of trout (*S. trutta*). However, we found galaxiids were a stronger regulator of individual growth than trout (H2b), which is consistent with ecological theory that predicts conspecific competition is stronger than interspecific competition for coexisting species (Adler et al., 2018; Chesson, 2000).

These results build on Howard’s (2007) findings where, under stable flow conditions, galaxiids grew slower within trout-free treatments than those with trout, suggesting that trout may indirectly enhance growth by reducing conspecific competition among galaxiids under otherwise crowded conditions. Thus, galaxiid growth rate is the result of combined con- and interspecific density-dependent mechanisms, which are punctuated by flood impacts. Density-dependent mechanisms like these are important for regulating population biomass relative to available resources (Watson et al., 2022).

Smaller-bodied juvenile galaxiids in low conspecific densities were most affected by the presence of trout in terms of growth. Although large-bodied trout (>150 mm) can consume non-diadromous galaxiids of all sizes (McIntosh, 2000), juvenile galaxiids are particularly vulnerable to predation and competition due to their small body size and weak competitive ability (Jellyman et al., 2017), and likely less capable of evasive maneuvers compared to adults (David & Closs, 2002). Consequently, stable streams supporting high abundances of trout often have poor representation of small size classes of galaxiids (McIntosh et al., 2010). This mechanism is represented in streams globally where introduced fish compete with natives. For example, the presence of introduced mosquitofish (*Gambusia affinis*) substantially reduced both the survivorship and individual growth of juvenile least chub (*Iotichthys phlegethonti*), suggesting the effects of introduced fish competition was to reduce native growth and prolong the period of size-selective predation (Mills et al., 2004). We expect juvenile galaxiids to experience the same interplay of reduced juvenile survivorship and growth in the presence of trout, which could also prolong their vulnerability to size-selective flood mortality.

### Conservation implications

Our results together present evidence for a combination of extreme flood disturbances and introduced trout affecting native non-diadromous galaxiid population dynamics. Although the threats against native fauna are diverse, many conservation tools are available to assist native fish population persistence. For instance, maintaining strategically placed trout-free source populations of adult galaxiids could be an important tool to enable native dispersal into trout-affected habitat and maintain population resilience, given that recruitment of non-diadromous galaxiids is higher in the absence of trout (Woodford & McIntosh, 2010). In extreme cases, maintaining stable sections of deep pools and backwaters within flood prone rivers could be an effective tool to provide refuge for adult native fish during high flow events (McIntosh & Barrett, 2022; Sedell et al., 1990), which can recolonise vacant reaches after disturbance (Davey & Kelly, 2007). However, in-stream refuges that seek to benefit native galaxiids could also promote the presence of trout and increase pressure on natives, given salmonids also prefer pools for the deep-water refuge and cooler water they provide during droughts (Torgersen et al., 1999; Young et al., 2010). Managers should therefore consider the indirect effects of conservation action in trout invaded habitats and prioritise refuge maintenance for native galaxiids within trout-free reaches to avoid indirectly exacerbating the pressure of introduced trout on vulnerable native species.

Individual growth of native galaxiids was dependent on the density of both trout and galaxiids. Extreme floods that reduce abundances of all size classes could, therefore, release non-diadromous galaxiids from inter- and conspecific competition, as well as trout predation, and promote growth of the few remaining individuals through density-dependent mechanisms. As a result, galaxiids that persist through extreme flood events, particularly resistant adults, and experience enhanced individual growth from reduced competition (Ali et al., 2003), could compensate for low abundances of galaxiids after extreme flood events by increasing maturation rates and egg production (Letcher & Terrick, 1998). However, compensatory responses within native galaxiid populations may be overshadowed by introduced trout reducing density-dependent growth that, in the absence of trout, would otherwise promote population recovery. Thus, the presence of trout in highly disturbed systems, perhaps driven by reinvasion from stable source populations nearby, may impair galaxiid recovery and exacerbate the threat of future floods on populations of native galaxiids. Taken together, these results highlight the need to prioritise the protection of vulnerable native fish populations from non-native species in catchments susceptible to increasing frequencies and magnitudes of floods.

The predicted increased frequency of extreme hydroclimatic disturbance events (IPCC, 2021) suggests that disturbances will have an increasingly common role in regulating native species in the future. Yet, we are currently ill-prepared to respond to such ‘black swan’ events given the lack of empirical research on their impacts (Macinnis-Ng et al., 2024). If extreme events occur more frequently than the period required for population recovery, then population dynamics of native species may remain in a constant state of unstable recovery and face an increased risk of local extirpation, particularly with the background influence on non-native competitors. This is indeed possible, with recovery of populations and communities from extreme hydroclimatic events being highly context dependent and, in certain circumstances, not possible (Tonkin, 2022). Indeed, past research has shown that riverine species richness, community composition and fish density can take up to 3 years to reach a stable state after a human induced ‘fish kill’ event (Rohr et al., 2021). In Australia, research has shown that galaxiid species may require a similar period of recovery, where *G. olidus* took 3 years to reach stable breeding populations after trout eradication (Lintermans, 2001). However, long term studies of riverine fish population dynamics are rare, especially in New Zealand (Hayes et al., 2010, 2019). Therefore, longer-term research is needed to gain a better understanding of non-diadromous galaxiid population dynamics in increasingly dynamic environments and in the presence of non-native trout. By monitoring population recovery over multiple seasons (i.e., >3 years), the dynamics of populations, including various vital rates of species, can be captured more fully (Verhelst et al., 2022). Furthermore, using multiple measures of population vital rates, such as native population growth, recruitment and individual growth can provide a well-rounded mechanistic assessment of native recovery dynamics, which holds one of the keys to developing effective management tools that preserve native freshwater fauna in a rapidly changing climate.

## Author contributions

R. Lennox, A. McIntosh, J. Tonkin and N. Boddy conceived the ideas and designed methodology; R. Lennox, J. Tonkin, N. Boddy and H.R. Lai collected the data; R. Lennox, H.R. Lai, D. Stouffer and C. Zammit analysed the data; R. Lennox led the writing of the manuscript. All authors contributed critically to the drafts and gave final approval for publication.

## Acknowledgements

We thank the many landowners that provided us access to some of the sampled streams. We also thank members of the Freshwater Ecology Research Group and Tonkin Lab at the University of Canterbury for feedback on project development, and Olivia Hore for providing data for six of the twelve sites from sampling prior to the flood event. We also thank the many dedicated assistants and volunteers who contributed to the 2021–2022 fieldwork. We thank MetService NZ Ltd. for supplying rainfall data, and Justin Rogers’ assistance with handling this data. Funding was provided by the Department of Conservation (NOF-BIO-263). JDT is supported by a Rutherford Discovery Fellowship administered by the Royal Society Te Apārangi (RDF-18-UOC-007), and Te Pūnaha Matatini, a Centre of Research Excellence funded by the Tertiary Education Commission, New Zealand. HRL was supported by the Marsden Fund Council administered by the Royal Society Te Apārangi (MFP-UOC2102). JDT and ARM acknowledge funding from the Ministry of Business, Innovation and Employment (Fish Futures: preparing for novel freshwater ecosystems; CAWX2101). JDT and HRL acknowledge funding from Bioprotection Aotearoa.

## Data availability statement

Data will be archived on Dryad

## SUPPORTING INFORMATION

### Appendix 1: Additional details on methods

#### Estimation of annual recurrence intervals using the New Zealand Convective Scale Model

The hydrological simulations were generated using the default surface water model used with the New Zealand Water Modelling Framework (NZWaM), TopNet (Clark et al., 2008). TopNet is a spatially distributed, hourly time stepping model that combines conceptual water balance models for each sub-catchment (Beven et al., 1995) with a kinematic channel routing model (Goring, 1994) to route streamflow to the basin outlet. TopNet was designed to have a sufficiently comprehensive description of catchment hydrology to be used in the diverse range of hydro-climate landscapes present in New Zealand (McMillan et al., 2016). To maintain consistency in both gauged and ungauged catchments, TopNet is uncalibrated to observed discharges and uses nationally available model parameters (Cattoën et al., 2022). It is widely used in hydrological modelling applications in New Zealand at catchment scale or national scale ranging from operational flow forecasting (Cattoën et al., 2022; McMillan et al., 2013), to predict the hydrological impacts of climate change (Collins, 2020; Poyck et al., 2011), and for national water accounting (Griffiths et al., 2021).

As part of this application, TopNet has been coupled with the New Zealand Convective Scale Model (NZCSM), a local implementation of the UK Met Office Unified Model (Bush et al., 2020). NZCSM is a convection permitting deterministic model, with a grid resolution of 1.5 km available since July 2015. Hourly time series discharges were generated at each location of interest over the period of simulation (2015-2022). A flood frequency analysis was completed, for each site, on the annual maximum daily average peak discharge implemented with the *extRemes* R package (Gilleland & Katz, 2016; R Core Team, 2022). The analysis assumes that the General Extreme value distribution follows a Gumbel distribution (Henderson et al., 2018) and a return period was hence derived.

#### Von Bertalanffy growth function

The von Bertalanffy growth function is most commonly written as

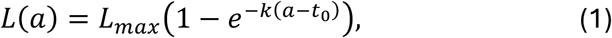

where *L*(*a*) is fish length *L* at age *a*, *L*_max_ is the maximum achievable length, *k* is the growth coefficient, and *t*_0_ is the theoretical age at which *L*(*t*_0_) = 0. This expression arises as a direct result of the less commonly written linear differential equation

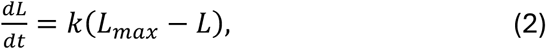

which specifies the *instantaneous* growth rate of an individual fish of length *L* based on the same parameter values. Given the theoretical initial size *L*(*t*_0_) = 0, corresponding theoretical age *t*_0_, and target age *a*, the solution to the initial value problem

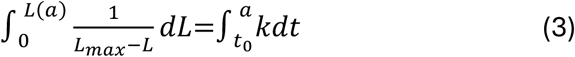

is exactly equal to Eq. (1).

Alternatively, one can use the von Bertalanffy growth model (Eq. 2) to predict a fish’s future size at a future time *L*(*t*_1_) given knowledge of its current size *L*(*t*_0_) and how much time has elapsed (i.e., *t*_1_ − *t*_0_). This is similarly achieved by integrating the dynamical equation

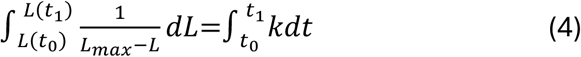

and solving for the single unknown, *L*(*t*_1_). This solution is given by

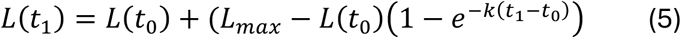

#### Specification of priors used in Bayesian inference

For all models, we used weakly informative priors that were default in the *brms* package, except for the following parameters to facilitate model convergence and chain mixing. For both count and growth models, we imposed a stronger prior of Student-*t*(1, 0, 0.5) for the standard deviations of all random intercepts. For the growth model, we further imposed a strong prior for *k*_0,s_ using Normal(−7, 0.5) and for *L*_max,s_ using Normal(5, 0.5); these values were determined by examining the distribution of the observed growth and maximum length.

## Supporting figures

**Figure S1:**
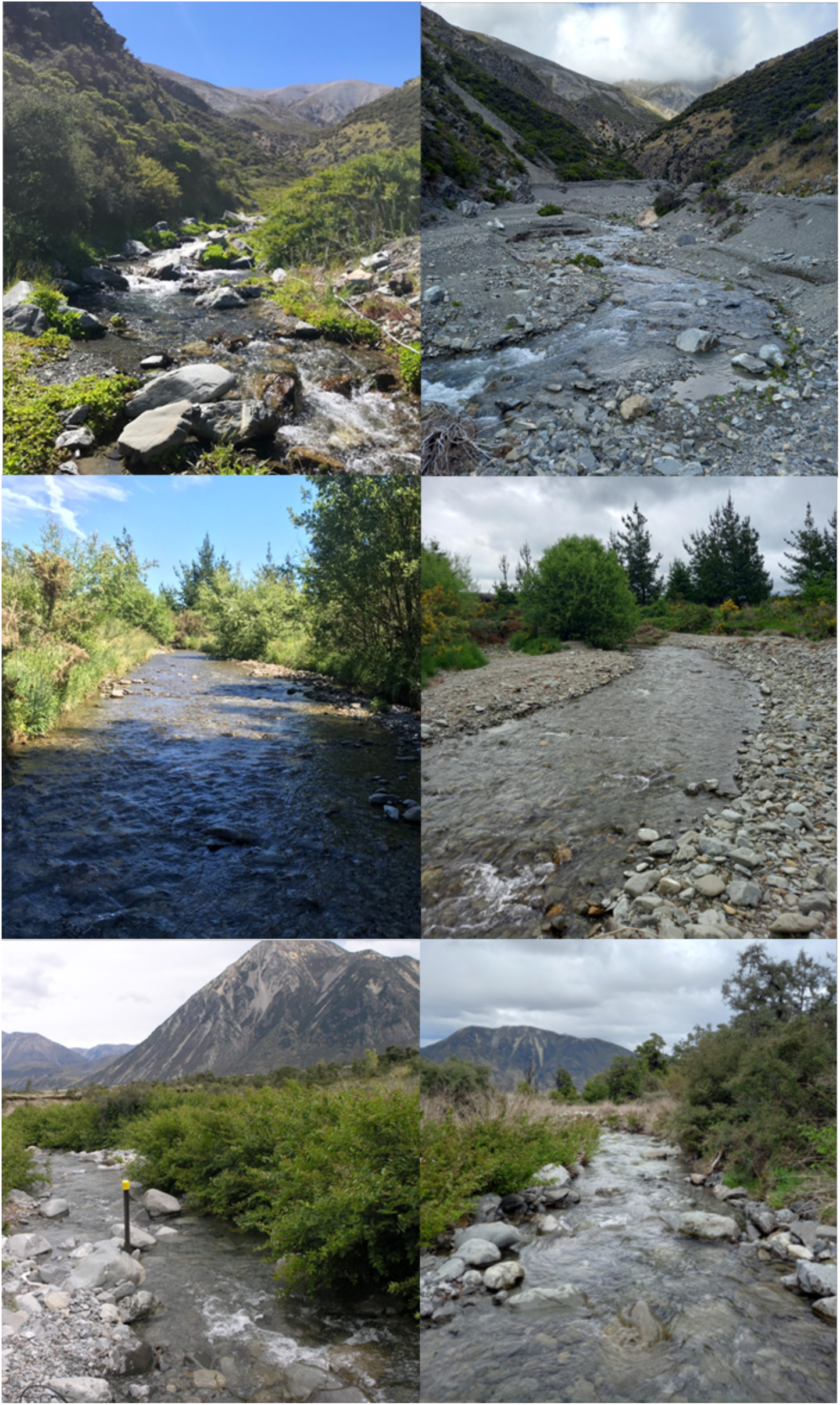
Photos of three sites, arranged in ascending order of flood magnitude, sampled throughout the Canterbury High Country and foothills in the 2021-21 season before (left) and 2021-22 season after (right) the 2021 Canterbury flood event. Top; Dry Stream (estimated 70-year ARI). Middle; Little Kowai (estimated 30-year ARI). Bottom; Lower Farm (estimated 5-year ARI). Photos before the flood event were taken by O. Hore, and photos after the flood were taken by R. Lennox.

**Figure S2:**
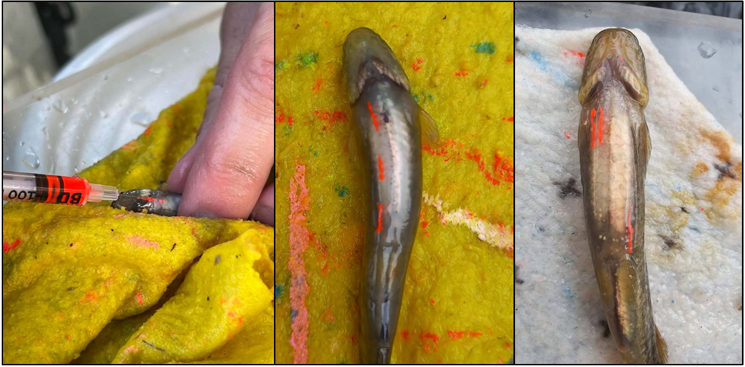
Tagging process using a Visual implant elastomer (VIE), demonstrating the placement of tags along the ventral side to create a unique identity for two individual adult non-diadromous galaxiids. The orange markings indicate both fish were first caught and tagged in the first sampling round. Left, tagging using a hypodermic needle. Centre; tag ID 1O2O3O. Right, tag ID 1O1O6O. Photos: R. Lennox.

## Supporting tables

**Table S1:**
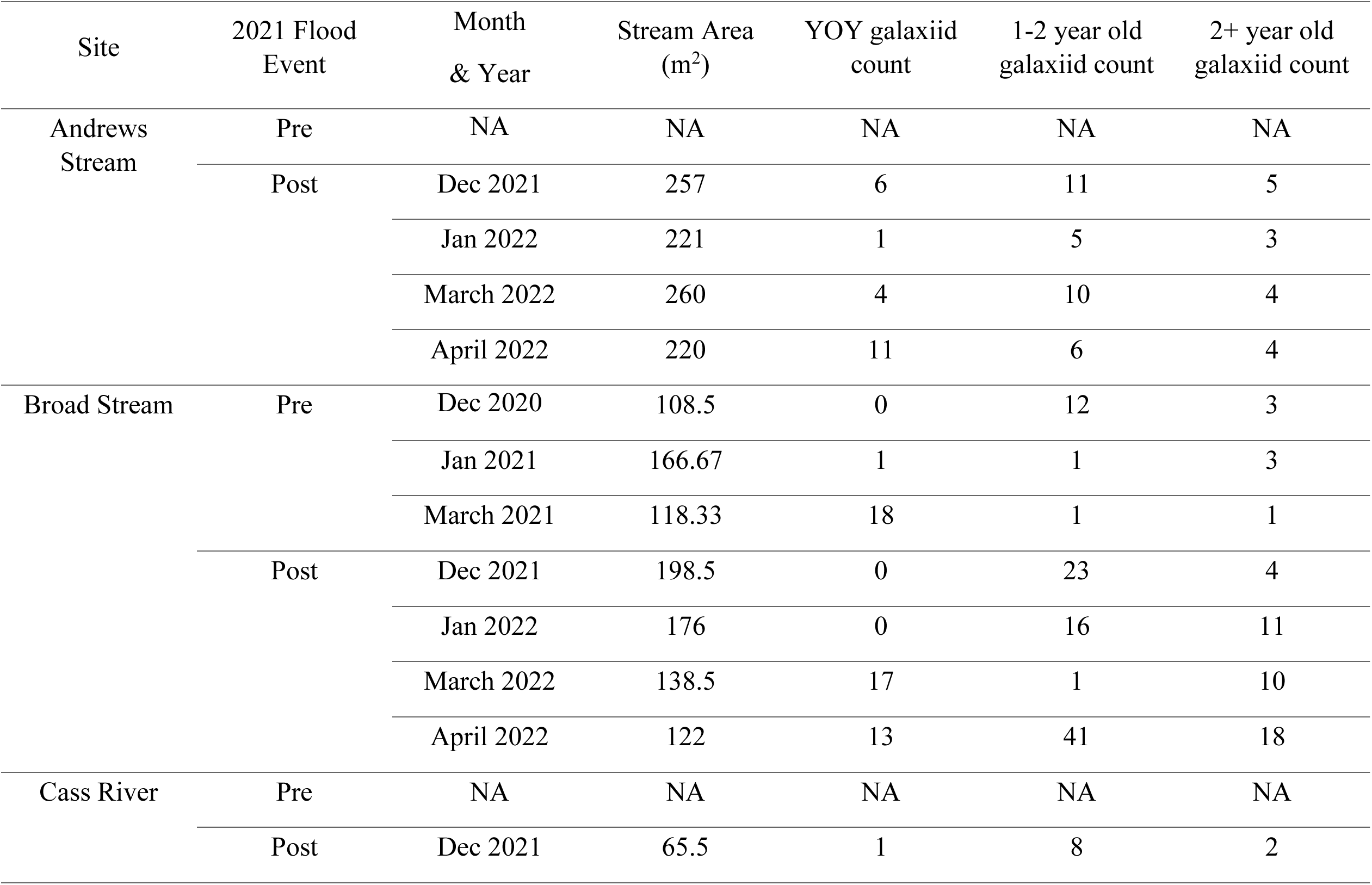
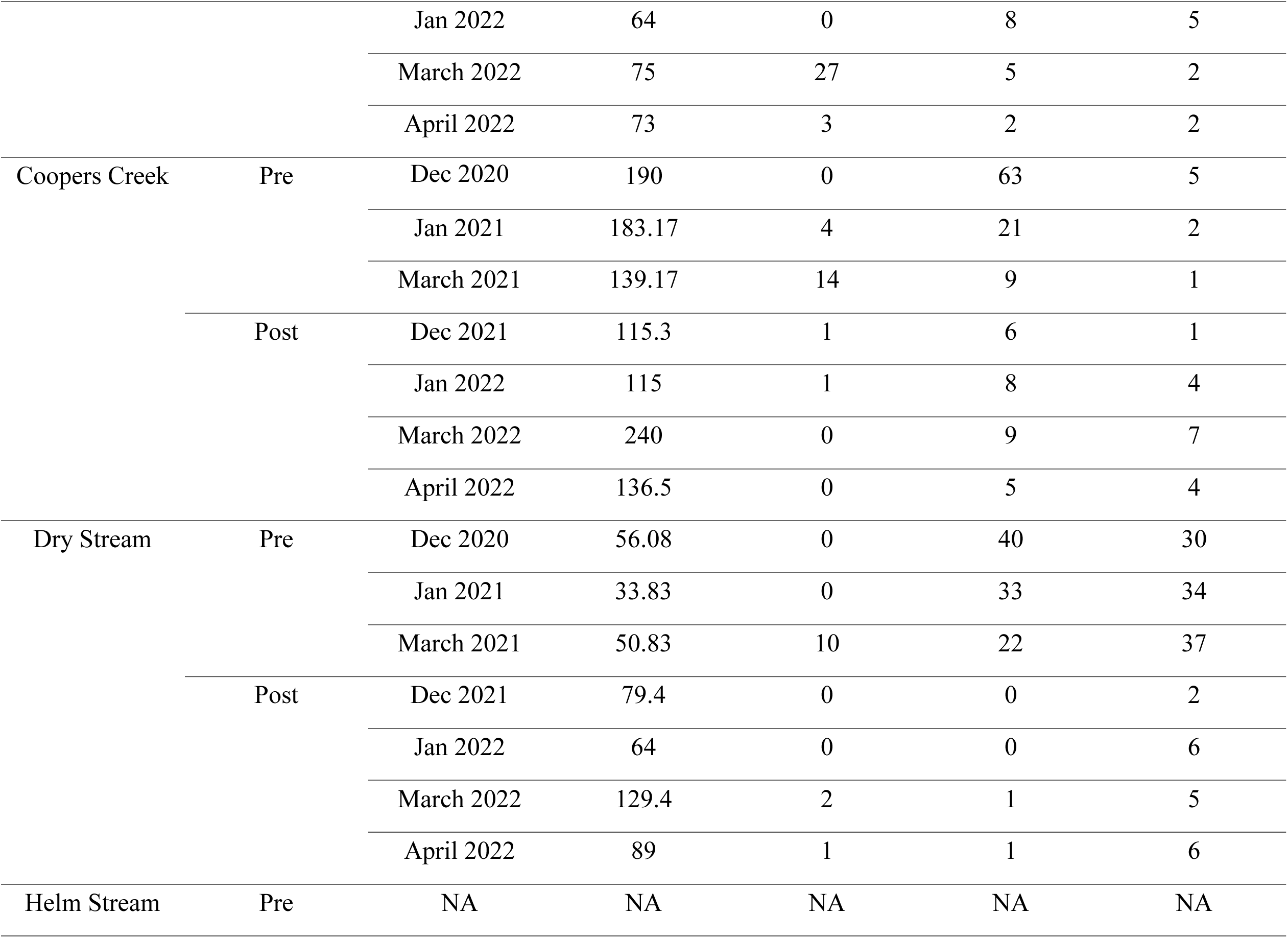
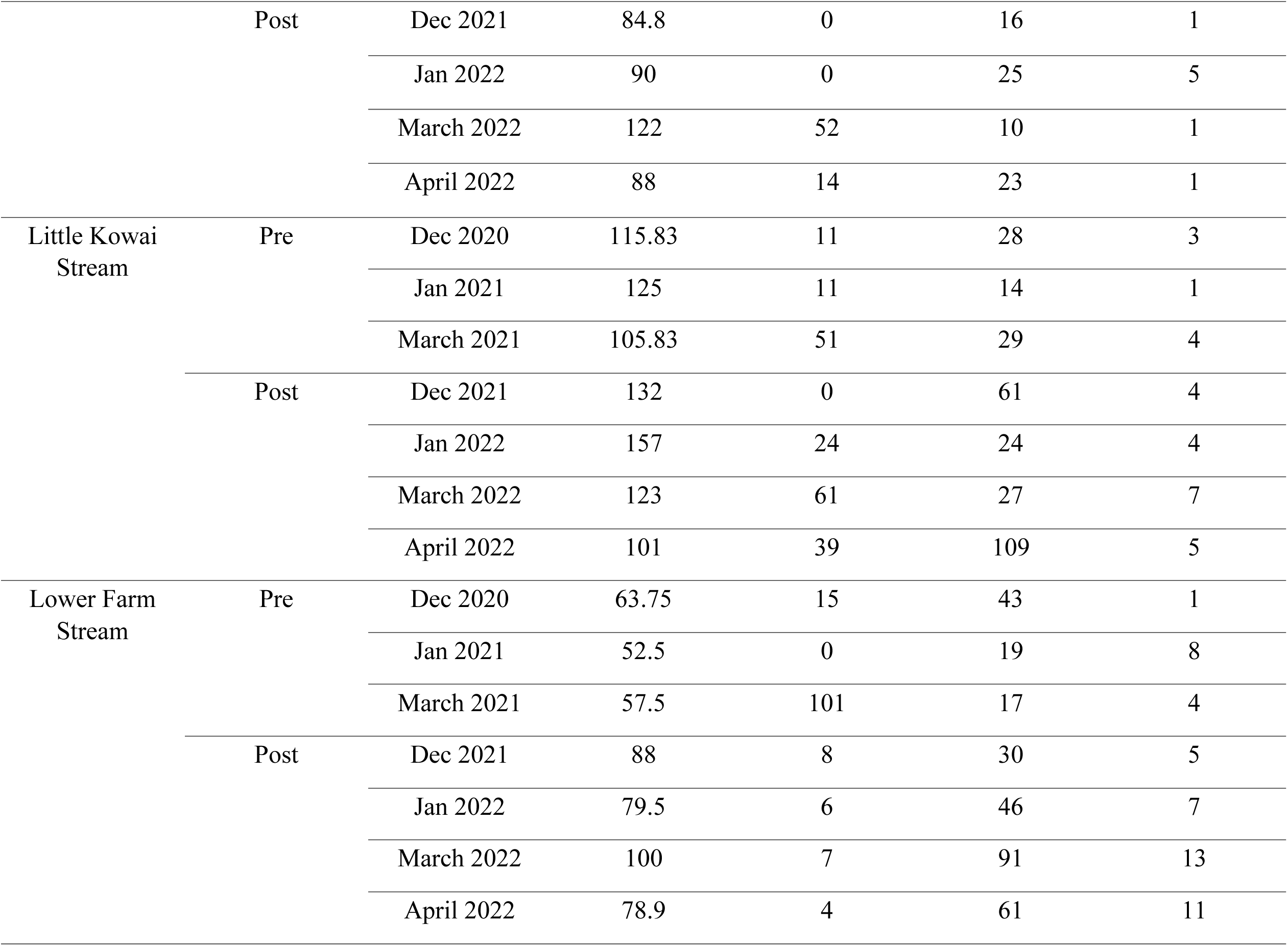
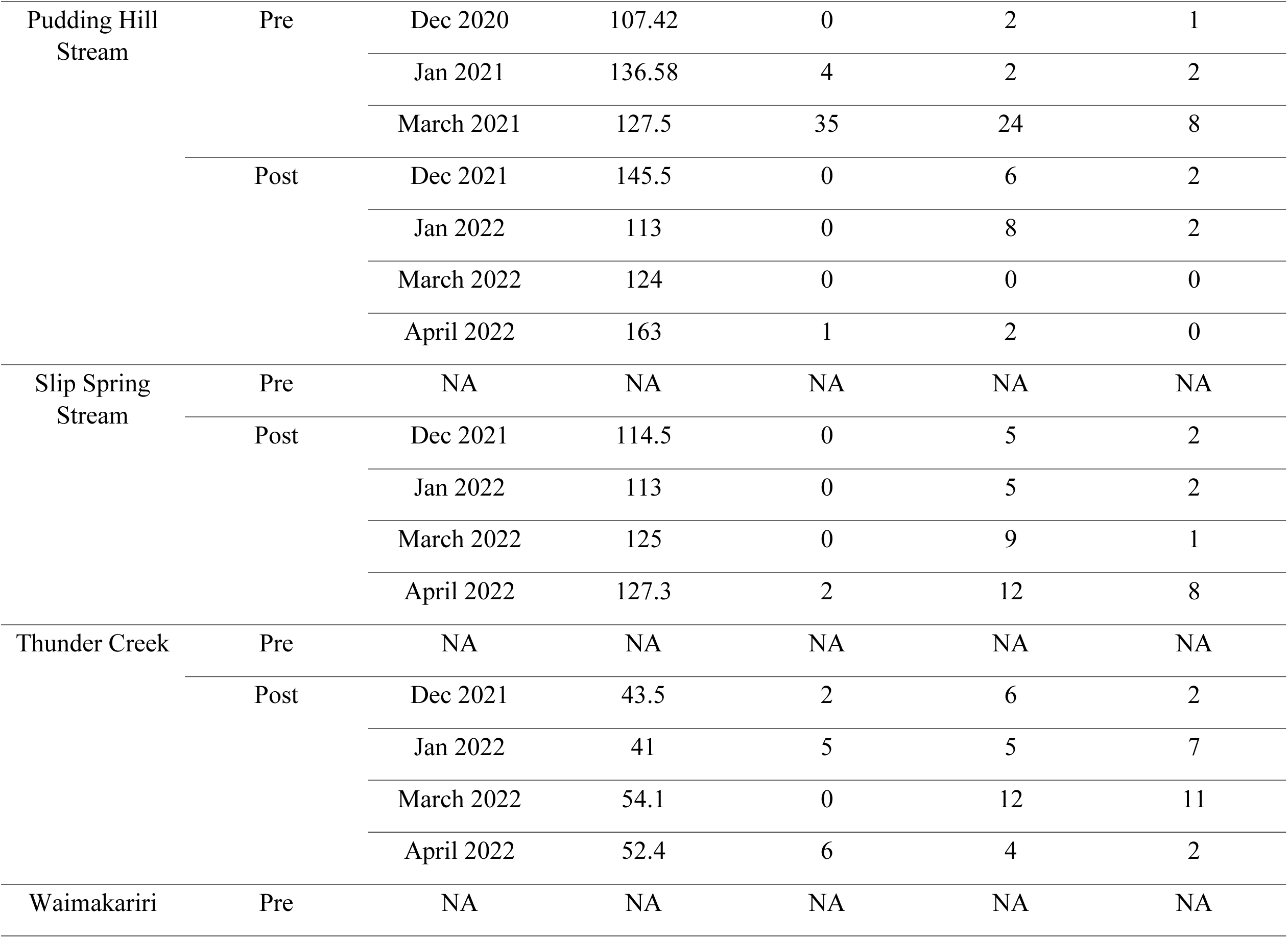
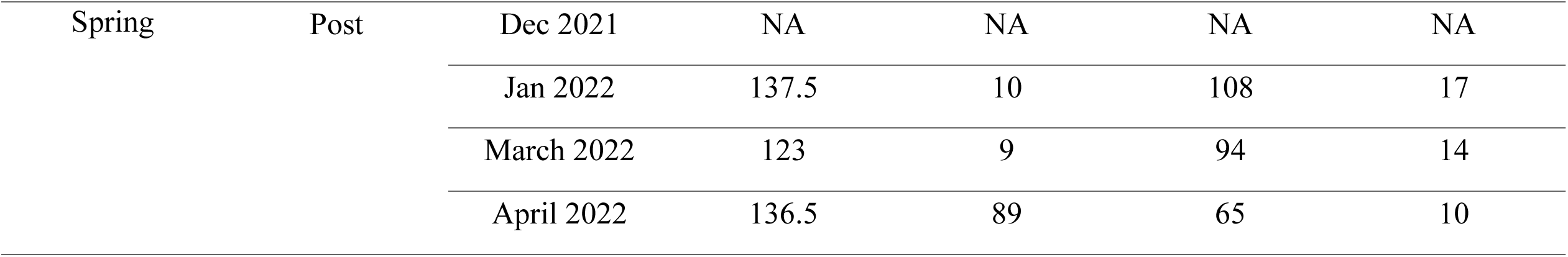
Twelve streams in the Canterbury High country and foothills sampled before (2020-2021 summer season) and after (2021-2022) the 2021 Canterbury flood event.

**Table S2:** Coordinates (NZGD), 2021 Canterbury flood event magnitude (annual recurrence intervals) and trout abundance across the twelve sites sampled on four occasions during the 2021-2022 summer season. Asterisks denote sites with both pre- and post-flood samplings.

| Site | Latitude<br>(NZGD) | Longitude<br>(NZGD) | 2021 Flood<br>Magnitude<br>(ARI) | Trout<br>abundance |
| --- | --- | --- | --- | --- |
| Andrews Stream | -42.9930230S | 171.7930993E | 2-year | Common |
| Broad Stream* | -43.0354816S | 171.6515363E | 20-year | Absent |
| Cass River | -43.0257364S | 171.7482350E | 5-year | Rare |
| Coopers Creek* | -43.2715905S | 172.0954908E | 60-year | Rare |
| Dry Stream* | -43.2612272S | 171.7244908E | 70-year | Absent |
| Helm Stream | -43.3343168S | 171.6698248E | 70-year | Common |
| Little Kowai Stream* | -43.3023396S | 171.9062573E | 30-year | Rare |
| Lower Farm Stream* | -43.0014570S | 171.8092362E | 5-year | Absent |
| Pudding Hill Stream* | -43.6061190S | 171.5407277E | 60-year | Common |
| Slip Spring Stream | -43.25737481S | 171.7098206E | 40-year | Common |
| Thunder Creek | -43.1961746S | 171.7175645E | 20-year | Absent |
| Waimakariri Spring | -43.01614294S | 171.8119206E | 5-year | Rare |

